# Spef1 is a microvillar component that limits apical actomyosin contractility and preserves intestinal barrier function

**DOI:** 10.1101/2025.07.17.665359

**Authors:** Rocio Tapia, Angelo Morales, Gail A. Hecht, Matthew J. Tyska

## Abstract

Epithelial sheet integrity is established by adherent contacts that form between cells, at the interface between their apical and basolateral domains. Although cell contacts are reinforced by actomyosin contractility, which generates tension that propagates across the apical surface. How epithelial cells tune tension to reinforce junctions without compromising their physical barrier properties remains unclear. Herein, we report that Sperm Flagellar 1 (Spef1) is a microvillar component enriched in the apical domain and terminal web of enterocytes that prevents actomyosin hypercontractility. Loss-of-function in Caco-2 BBE cells showed that Spef1 depletion induced invaginations of the apical domain at tricellular contacts, with a redistribution of the tricellular and tight junction components. These changes, which were paralleled by increased activity of NM2A across the apical surface, compromised intestinal barrier function. These findings highlight Spef1 as a microvillar resident that tunes actomyosin contractility across the apical surface, to a level appropriate for junctional reinforcement and maintenance of epithelial function.

## INTRODUCTION

Epithelial monolayers play central roles in tissue compartmentalization, physically separating and regulating transport between physiologically distinct spaces. The cell-cell contacts that drive formation of these sheets are composed of several adhesion complexes that stratify along the apical-basolateral axis including tight junctions (TJs), adherens junctions (AJs), and desmosomes ^1^. These adhesive complexes are also rich in actin filaments and non-muscle myosin-2 (NM2), which generate the mechanical force required for supporting junctional integrity and regulating junctional permeability ^2^.

The establishment of apicobasal polarity defines two surface domains that are functionally distinct ^3^. Here, a major morphological distinction between the two arises from the assembly of apical specializations, which are typically comprised of one or more membrane protrusions that extend outward enabling interactions with the luminal environment ^3^. Depending on the tissue, these structures are supported by elaborate microtubule- or actin-based cytoskeletal assemblies ^4–7^. For example, the primary cilium is supported by axonemal microtubules (MTs) and extends from the apical surface of kidney tubule cells to sense the flow of filtrate ^4^. Similarly, unipolar actin bundle-supported brush border microvilli (MV) form on the apical surface of nutrient absorbing enterocytes to increase solute uptake potential ^5–7^.

The cytoskeletal architectures that support surface protrusions are conventionally viewed as separate from the actomyosin contractile networks associated with cell junctions. However, previous reports have shown that in the context of the intestinal brush border, these cytoskeletal features are closely connected to the TJs and AJs, suggesting that these networks may be physically linked ^8–14^. Indeed, super-resolution localization of NM2C in enterocytes revealed localization throughout a sub-apical plane that was continuous with the high-level enrichment in junctional belts ^15^. From this perspective, molecules that reside in brush border MV could play a role mediating cell contractility. In this work, we identify Sperm Flagellar 1 (Spef1, ∼25 kDa) as one such molecule.

Spef1, also known as CLAMP (CaLponin-homology And Microtubule-associated Protein), was previously identified in the flagellum of spermatids and in the cilium ^16, 17^. In these contexts, Spef1 associates with MTs, promotes bundle formation and stabilizes MTs against cold-induced depolymerization ^16, 18–22^. Structural analysis of the tip-central pair (CP) of *Tetrahymena*, revealed that Spef1 makes electrostatic contacts with α- and β-tubulin, to function as a seam-binding protein that stabilizes and crosslinks tip-CP MTs (Legal et al., 2025, BioRxiv unpublished data). Spef1 localizes along MTs bundles of pillar cells in the organ of Corti, and distributes as puncta along the CP apparatus in mature cilia where it participates in the CP formation and regulates the rotational beat of the ependymal cilia ^18, 20, 23, 24^.

Despite its localization in the cilia tips, Spef1 is also enriched at the apical surface and at the cell-cell contacts of the multi-ciliated epithelium ^19, 25^. The apical positioning of Spef1 has been explored during the cell intercalation in *Xenopus leavis* embryo where it promotes MTs stabilization, resulting in an increased tubulin acetylation along the axis of cell migration. As migrating cells reach the outer epithelia, Spef1 is observed at the leading edge of the intercalating cells, partially colocalizing with components of the Par polarity complex such as αPKC, and this interaction is crucial for the directed cell migration ^19^. At cell edges, Spef1 contributes to the asymmetric distribution of membrane-associated planar cell polarity (PCP) proteins in multi-ciliated epithelia, suggesting that Spef1 mediates cell migration and PCP signaling pathways ^25^.

Recently, our group demonstrated that Spef1 is present at the apical surface, cell-cell contacts and basolateral domain of intestinal epithelial cells ^26^. This pattern of localization depends on the degree of cell polarization. In sparse, migrating intestinal epithelia, Spef1 is enriched at the basal membrane in stress fibers and focal adhesion sites. When epithelia reach full polarization and cell migration is reduced, Spef1 is localized at cell contacts and in the apical domain ^26^. In a wound healing model of intestinal epithelia, Spef1 is accumulated at the edges of migrating cells ^26^. Loss of Spef1 disrupted filopodia and lamellipodia assembly, with disorganization of actin filaments and misoriented filopodia ^26^. Although there is no evidence on the direct binding of Spef1 to actin, its full-length sequence which consists of a NH_2_ -calponin homology (CH) region and a coiled-coil (CC) -COOH domain shares homology with numerous cytoskeletal proteins ^26^. Additionally, Spef1 forms a complex with adhesion molecules that interact with the actin cytoskeleton ^26^. Although, collectively, these prior studies suggest actin-based functions for Spef1, the role of this factor in the apical domain of the transporting epithelia remains unclear.

In this paper, we explore the role of Spef1 in the context of the intestinal epithelium, specifically at the enterocyte apical surface, which lacks MTs and is home to the actin-rich brush border. Super-resolution imaging revealed that endogenous Spef1 is enriched in brush border MV, including the rootlet segment that extend down into the terminal web. Loss of Spef1 disrupted the organization of epithelial monolayers causing major defects in tricellular and bicellular contacts. These morphological changes were accompanied by the failure of the establishment of the intestinal barrier function. Importantly, Spef1 depletion resulted in an increased localization of NM2A to the cell edges with an augmented actomyosin contractility. Inhibition of NM2 contractility rescued the Spef1 depletion phenotype. Together, our discoveries suggest that Spef1 is a microvillar and terminal web resident that controls actomyosin contractility across the apical surface, with implications for maintaining junctional architecture and epithelial function.

## RESULTS

### Spef1 is a resident of brush border microvilli

As a first step toward defining the function of Spef1 in the intestinal epithelium, we examined the localization of endogenous Spef1 using immunofluorescence (IF). We immunostained paraffin sections of mouse small intestine with probes for Spef1 and villin, an actin bundler and marker for the apex of the enterocyte. In agreement with previous studies ^26^, Spef1 was highly enriched in the apical domain of cells along the crypt-villus axis and demonstrated strong colocalization with villin (Fig. 1a). Additionally, fluorescence intensity quantification showed that apical Spef1 levels were significantly higher on the villus than in the crypt compartment (Fig. 1c). High magnification views of villus and crypt sections also revealed that Spef1 exhibited some basolateral localization in both regions (Fig. 1b,d). Interestingly, the lateral localization of Spef1 shifted from basal to apical as cells move out of the crypt and onto the villus (Fig. 1e). These results indicate that Spef1 levels and localization were impacted by the degree of the intestinal epithelial cell differentiation, with the highest levels observed in the apical domain of villus enterocytes.

**Fig. 1:**
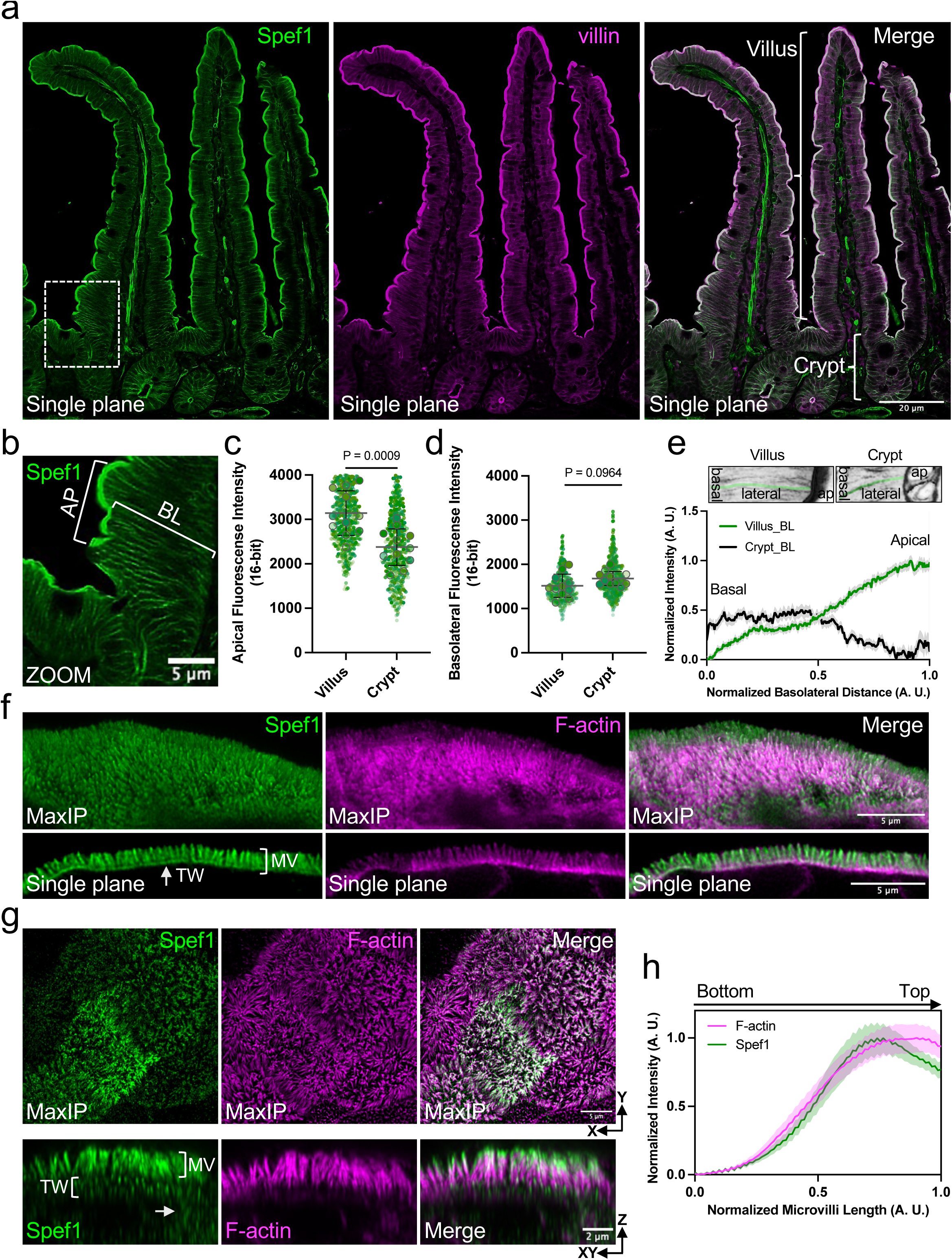
Spef1 localizes throughout the apical domain in differentiated intestinal epithelial cells. **a,** Single-plane confocal microscopy acquired from a mouse small intestinal tissue section oriented parallel to the crypt-villus axis; signals are anti-Spef1 (green) and anti-villin (magenta). Representative villus and crypt are highlighted by brackets in the merge panel. **b,** Magnified view of Spef1 staining corresponding to the region in **a** (dotted box) showing differential enrichment at the apical (AP) and basolateral (BL) domains. **c,d,** Plots show average fluorescence intensities of Spef1 from the AP and BL domains in villus and crypt regions; **c** includes quantification of 578 cells from 11 villus (*n* = 11) and 531 cells from 11 crypts (*n* = 11); **d** includes quantification of 419 villus cells and 560 crypt cells. **e,** Quantification of Spef1 fluorescence intensity along the BL domain from cells in villus and crypt compartments (*n* = 317 villus cells; *n* = 246 crypt cells). **f,** Maximum intensity projection (MaxIP) images of small intestinal tissue visualized with NSPARC illustrates endogenous Spef1 (green) and F-actin (magenta) in MV region. Single-plane images revealed presence of Spef1 at the base of MV (MV) in the terminal web (TW) co-localizing with F-actin. **g,** Projected NSPARC images of Caco-2 BBE monolayers (21 DPC) showing the localization of endogenous Spef1 (green) and F-actin (magenta). **h,** Normalized fluorescence intensity of F-actin and Spef1 along the axis of MV (*-xz* sections). From three separate experiments (*n* = 3), the number of fields quantified = 15. *P* values were calculated analyzing the standard deviation (SD) using a two-tailed *t*-test.

Given its strong localization to the apex of the enterocyte, we next asked if Spef1 is a resident of brush border MV. To address this question, frozen tissue sections from mouse small intestine were stained for Spef1 and subject to super-resolution microscopy (Nikon SPatial ARray Confocal, NSPARC). Images from the villus revealed that Spef1 targeted to MV and showed a strong co-localization with F-actin (Fig. 1f, top). Lateral views of the brush border also revealed that Spef1 signal extended along the full length of MV core bundles, down to their basal rootlets (Fig. 1f, bottom, terminal web, TW, arrows). To further investigate Spef1 function, we employed Caco-2 BBE cells, which develop an enterocyte-like phenotype as they differentiate over the course of several weeks in culture ^27^. Caco-2 BBE cells were immunostained for endogenous Spef1 and co-stained with phalloidin to label F-actin at 21 days post-confluence (DPC). NSPARC imaging of these samples revealed that, similar to native intestinal tissues, Spef1 signal was strong in the apical domain, where it exhibited robust colocalization with the F-actin rich core bundles that support MV (Fig. 1g). Dimmer Spef1 puncta were also observed deeper in the cytoplasm (Fig. 1g, bottom arrow). Quantification of Spef1 and F-actin signals using line scans revealed highly correlated intensities along the full axis of MV (Fig. 1h). These findings provide further indication that Spef1 is a *bona fide* resident of the actin-rich brush border in differentiated intestinal epithelial cells; they also suggest that the Caco-2 BBE monolayers are a suitable model to investigate Spef1 function in this context.

### Spef1 is required for maintaining the integrity of the apical surface

To investigate the role of Spef1 at the apical surface, we generated Caco-2 BBE Spef1 knock-down (Spef1 KD) cells using an inducible (Tet-On) expression system. Spef1 depletion was induced with doxycycline (dox) and a time course-dependent loss of expression was confirmed using IF and western blot (WB) assays (Fig. S1). Using this model, we sought to determine how Spef1 depletion impacted the architecture and physiology of Caco-2 BBE cells. Control cells (7 and 21 DPC) exhibited a regular apical surface architecture (Fig. 2a,c, top). In contrast, Spef1 KD monolayers displayed marked changes in cell morphology (Fig. 2a,c, bottom). Segmentation analysis demonstrated that Spef1 KD cells exhibited apical surface areas that were highly variable, with a distribution that was shifted toward smaller values compared with control monolayers (Fig. 2b,d). In addition, the lateral edges of Spef1 KD cells (7 DPC) showed increased membrane tortuosity compared to control monolayers [Control = 1.1 ± 0.02 *vs* Spef1 KD = 2.17 ± 0.05] (Fig. 2e). Spef1 KD cells also developed increased cell heights compared with controls [Control = 11.3 ± 0.29 μm *vs* Spef1 KD = 20 ± 1.0 μm] (Fig. 2f). Whereas control Caco-2 BBE cells at 7 DPC exhibited a regular apical surface architecture and with accumulations of MV (Fig. 2g, top), Spef1 KD monolayers exhibited numerous aberrant F-actin-rich structures (Fig. 2g, bottom). Close inspection of *en face* (*xy*) and lateral (*xz*) sections of Spef1 KD cells revealed distinct classes of structures, including large invaginations that appeared continuous with the apical surface (<u>a</u>pical <u>m</u>embrane invagination<u>s</u>, AMIs) (Fig. 2h, top) and cyst-like features that were positioned more basally with no detectable contact with the luminal space (<u>b</u>asally <u>a</u>ccumulated <u>c</u>yst-like structure<u>s</u>, BACs) (Fig. 2h, bottom). Collectively, these data indicate that Spef1 plays a role in maintaining polarized intestinal epithelial cell architecture, and specifically, the integrity of the apical surface.

**Fig. 2:**
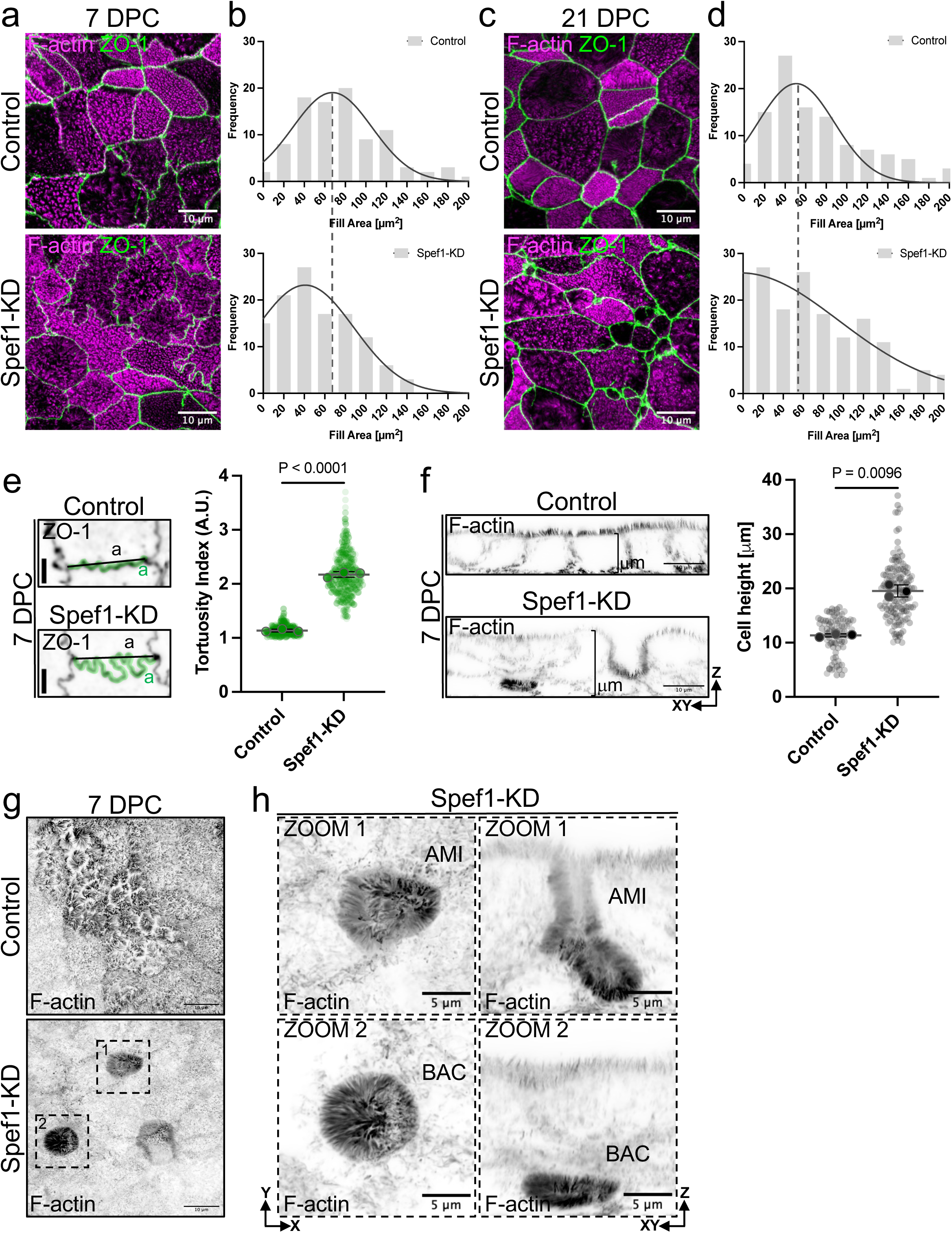
Spef1 depletion leads to a lateral expansion and basal accumulation of the apical compartment. **a,c,** Projected confocal volumes showing control and Spef1 KD cells 7 DPC (**a**) and 21 DPC (**c**) stained for F-actin (magenta) and ZO-1 (green) exhibit variations in the apical surface morphology in monolayers lacking Spef1. **b,d,** Frequency diagram of segmented cells from control and Spef1 KD condition at 7 and 21 DPC indicating the variability in the cell surface area during cell polarization. Nonlinear Gaussian curve was added to the histogram. For **b** and **d,** a total number of cells from several monolayers were segmented. For **b,** control = 94 cells, Spef1 KD = 118 cells; for **d,** control = 114 cells, Spef1 KD = 166 cells. **e,** Representative confocal *en face* images from Caco-2 BBE cells 7 DPC stained for ZO-1 (inverted channel) showing the AP domain. Plot represents the membrane tortuosity index (ratio a/a) of the apical surface. From three independent experiments (*n* = 3), control = 94 cells and Spef1 KD = 118 cells were quantified. **f,** Representative lateral confocal images from Caco-2 BBE cells 7 DPC stained for F-actin (inverted channel) showing the cell height. Plot represents the average of cell height values in the control and Spef1 KD condition (*n* = 3; with a total number of fields quantified for control = 84; Spef1 KD = 100). **g,** MaxIP confocal images from Caco-2 BBE cells 7 DPC stained for F-actin (inverted channel) showing the accumulation of MV structures. **h,** High magnification images of Spef1 KD cells from **g** (dashed boxes) illustrating extensive apical domain expansion and basolateral accumulation of these surface membrane structures. *P* values were calculated using two-tailed *t*-test.

### Spef1 KD monolayers lose AMIs and accumulate BACs over time

To enable detailed morphometry of the defects induced by Spef1 KD, we segmented confocal volumes based on the F-actin signal. The resulting 3D reconstructions allowed us to count and measure AMI and BAC structures (Fig. 3a). While such aberrations were occasionally observed in control Caco-2 BBE cultures, Spef1 KD cells developed a significantly higher number per field [Control_AMI_ = 1.15 ± 0.46 *vs* Spef1 KD_AMI_ = 13.81 ± 5.46; Control_BAC_ = 0.34 ± 0.74 *vs* Spef1 KD_BAC_ = 10.04 ± 4.13] (Fig. 3b,c). Previous studies demonstrated that Spef1 localization changes as epithelia acquire apical-basal polarity ^26^. We therefore investigated if the appearance of surface defects (AMIs and BACs) in Spef1 KD monolayers was dependent on cell polarization state. For these experiments, cells were cultured for 3 - 21 DPC and then fixed and stained with phalloidin to probe for F-actin. This enabled us to assess the number and size of AMIs and BACs at different time points (Fig. 3d-i). At the earliest time point (3 DPC), Spef1 KD monolayers contained large numbers of small AMIs per field [33.67 ± 3.26] (Fig. 3e), which were exclusively found at cell contacts (Fig. 3d). At 7 DPC, KD monolayers held fewer AMIs per field [9.0 ± 1.21] (Fig. 3e), but those that did form exhibited a striking increase in segmented volume (Fig. 3g). As monolayers matured, the number of AMIs per field further decreased and these structures were almost undetectable by 21 DPC (Fig. 3e-g). Interestingly, BAC appearance in differentiating Spef1 KD monolayers followed a contrasting trajectory, with number and segmented volume increasing significantly over time (Fig. 3e,h,i). These time-resolved findings suggest that Spef1 depletion leads initially to the appearance of AMIs, which preferentially form at cell contacts and likely represent precursors to BACs, which exclusively populate KD monolayers at later points in differentiation.

**Fig. 3:**
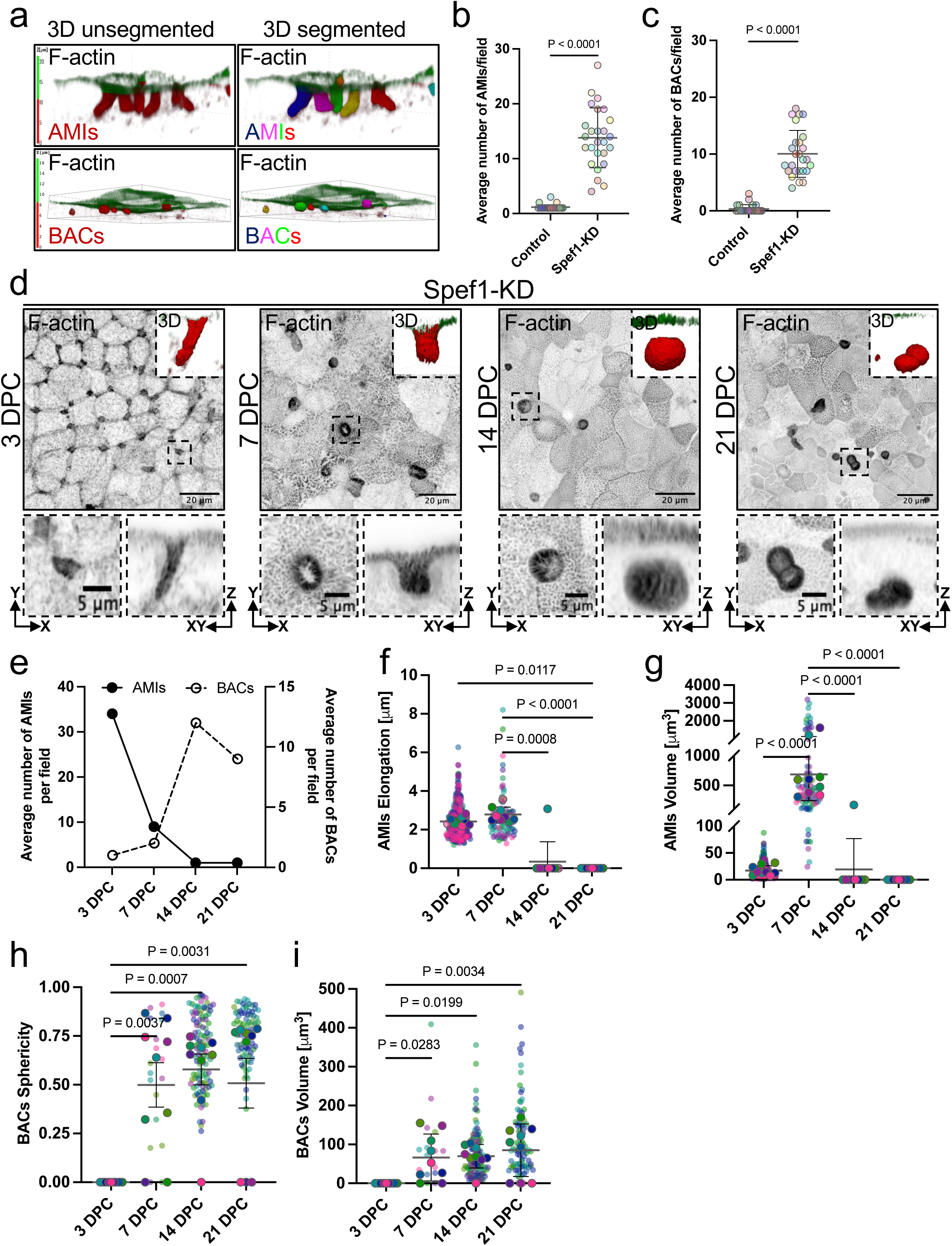
Loss of Spef1 leads to the formation of the aberrant MV structures. **a,** Representative 3D-rendered unsegmented (left) and segmented (right) images from Spef1 KD monolayers stained for F-actin (depth color coded) illustrating AMIs (top) and BACs (bottom). **b,c,** Plots indicate the average number of AMIs and BACs per field in control and Spef1 KD condition. The *n* values correspond to the total number of segmented fields per condition (*n* = 26). **d,** MaxIP confocal images of Spef1 KD cells (3 -21 DPC) labeled for F-actin (inverted channel). High magnification (bottom) regions *en face* and lateral sections illustrate AMI and BAC structures through cell polarization. **e,** Quantification of the average number of AMIs and BACs formed during cell polarization (3 - 21 DPC) in Spef1-depleted cells. **f**-**i,** Superplots represent the quantification of segmented AMIs and BACs through cell differentiation; elongation (**f**), volume (**g,i**) and sphericity (**h**) parameters were measured. For **e-i**, total number of fields (*n* = 9) were quantified in each condition. *P* values were calculated using two-tailed *t*-test.

### Spef1 is required for maintaining the integrity of tricellular and bicellular junctions

The staining results outlined above suggest that AMIs initially form at cell contacts, and potentially tricellular vertices. To test this idea, we stained control and Spef1 KD monolayers with tricellulin, a marker of the tricellular junctions (TCJs). Tricellulin was highly enriched at tricellular vertices and present to a lesser extent at the bicellular junctions at 7 DPC (Fig. 4a, arrowheads). In contrast, Spef1 KD cultures demonstrated much lower levels of tricellulin throughout the monolayer (Fig. 4b, left). Consistent with low levels of adhesion at these sites, Spef1 KD TCJs also demonstrated an aberrant “open” morphology, with villin extending down into these invaginations (Fig. 4b, zoom, *- xz* sections). As monolayers matured (21 DPC), the tricellular space eventually closed, although tricellulin signal remained aberrantly distributed along the lateral surface of the interface (data not shown). Collectively, these findings prompted us to hypothesize that Spef1 depletion leads to opening of tricellular contacts and expansion of the apical surface down the lateral surface of cells at these interfaces.

**Fig. 4:**
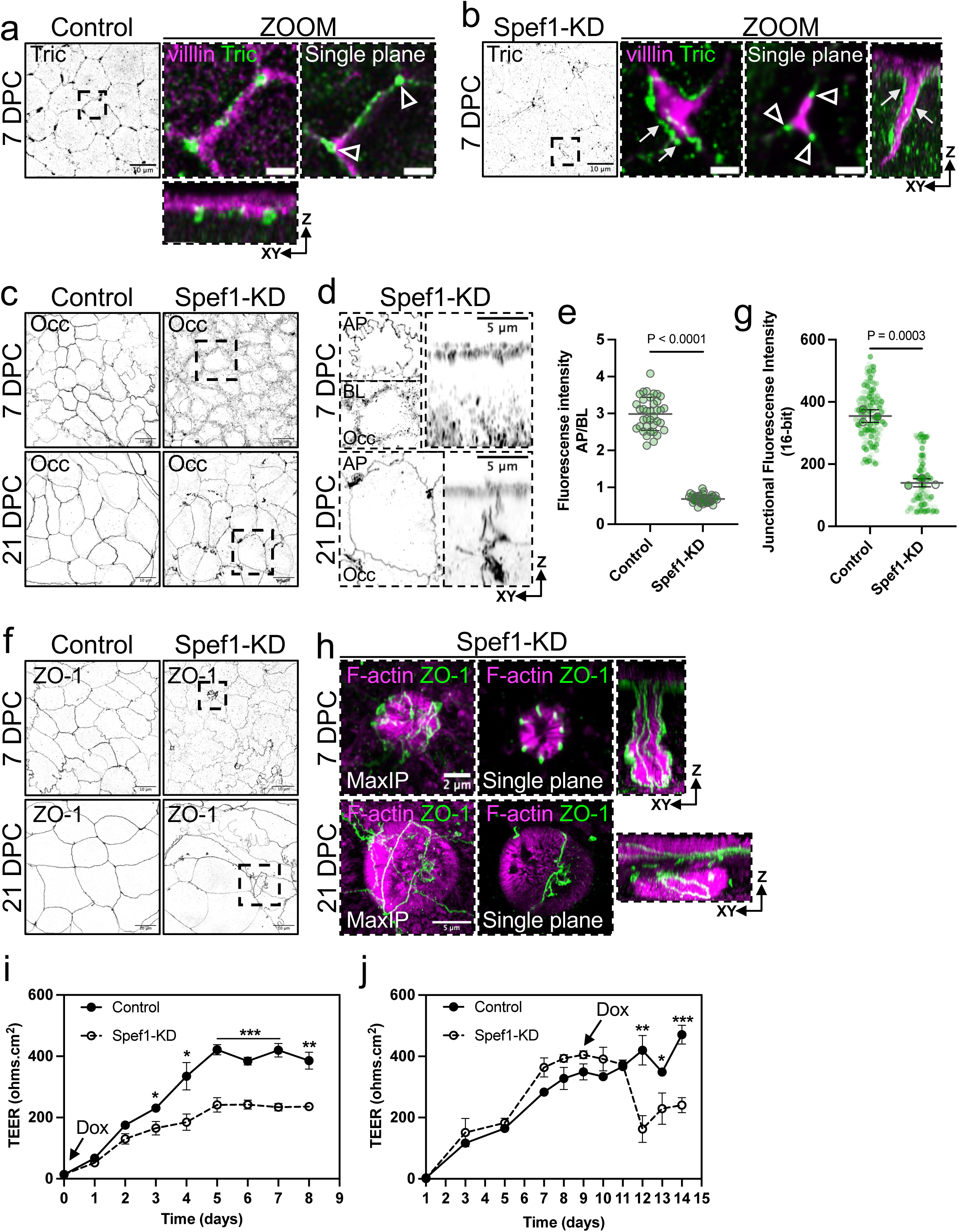
Spef1 depletion alters TCJ and TJ stability resulting in an impaired barrier function. **a,b,** Confocal images from control and Spef1 KD cells growing at 7 DPC immunostained for Tricellulin (Tric) and villin. High magnification showing top and lateral sections of merged images. Arrows and arrowheads indicate the localization of tricellulin along the lateral membrane or at the tricellular contacts, respectively. Scale bars, zoom: 2 μm. **c,d,** Confocal images from control and Spef1 KD cells plated at 7 and 21 DPC immunostained for occludin (Occ). High magnification (**d**), showing AP, BL and tricellular accumulation of occludin in the KD condition. **e,** Plot represents the ratio of occludin fluorescence intensity from the AP region involving the cell-cell contacts, and occludin staining accumulated at the BL membrane domain, bellow TJs of Caco-BB2 monolayers plated at 7 DPC. The *n* values correspond to the number of cells quantified (*n* = 50) per condition. **f,h,** Confocal images from control and Spef1 KD cells plated at 7 and 21 DPC immunostained for ZO-1 and F-actin. High magnification images indicate the localization of ZO-1 along the lateral domain and accumulated at the BACs (**h**). **g,** Plot represents the average of fluorescence intensity of junctional ZO-1 observed in control *vs* Spef1 KD cells grown at 7 DPC. The *n* values represent three independent experiments (*n* = 3) with a total number of quantified cells = 150 per condition. **i,j,** Tight junction assembly (**i**) and disassembly (**j**) was determined on SKCO-15-Spef1-KD Tet-On cells. For TJ assembly (**i**), induction with dox was achieved at D0, and for TJ disassembly (**j**) at D9. The TEER was measured every day. *P* values were calculated using two-tailed *t*-test analysis.

Based on previous studies, TCJ defects are likely to impact epithelial barrier function ^28–31^. To investigate potential barrier function defects in response to Spef1 depletion, we first explored the localization of the TJ proteins, occludin and ZO-1. As expected, control monolayers showed a well-defined pattern of occludin at the cell-cell contacts in undifferentiated as well as in fully polarized monolayers (Fig. 4c, left). In contrast, occludin was aberrantly localized at the basolateral membrane domain of cells in undifferentiated Spef1 KD monolayers (Fig. 4c,d, top). The portion of occludin fluorescence intensity at the junctional area *vs* the basolateral region was significantly lower in Spef1 KD (7 DPC) cells compared to control monolayers [Control = 2.9 ± 0.44 *vs* Spef1 KD = 0.68 ± 0.10] (Fig. 4e). Occludin was localized at the TJ region and was enriched at TCJs until the Spef1 KD cells reached a fully polarized state (Fig. 4c,d, bottom). Importantly, ZO-1, which contributes to TJ formation, apical surface architecture, and cortical F-actin network ^32–34^, was also impacted by Spef1 depletion (Fig. 4f). ZO-1 fluorescence intensity was significantly reduced at TJs in Spef1 KD monolayers (7 DPC) when compared to controls [Control = 355 ± 20 *vs* Spef1 KD = 140 ± 13] (Fig. 4g). Close inspection of Spef1 KD cells revealed that ZO-1 was laterally distributed where it appeared to enclose the apical membrane invaginations (Fig. 4h, top). High magnification images denote ZO-1 surrounding BAC structures from the top to the bottom side of the monolayers (Fig. 4h, bottom). These findings showed that Spef1 depletion mislocalized down into the lateral domain of neighboring invaginated cells, TJ components which appear enclosing BAC structures at the bottom of the polarized epithelia.

Given these striking defects in junctional markers, we next sought to determine how loss of Spef1 impacts the development and maintenance of transepithelial resistance (TEER), an established measurement for epithelial barrier function. For these experiments we generated a doxycycline-inducible Spef1 KD model using the human intestinal epithelial line, SKCO-15, which develops higher TEER values as they differentiate as opposed to the parent line Caco-2 BBE cells (21 DPC) that exhibit relatively low TEER (120 Ω.cm^2^). To monitor TEER during TJ assembly in SKCO-15 cells, control and Spef1 KD monolayers were incubated on Transwell filters with or without dox from day 0 (D0) to D8 (Fig. 4i). To explore whether Spef1 depletion affected the established barrier function, monolayers were grown without dox until they reached stable TEER values (D9), then dox was added to induce Spef1 KD (Fig. 4j). TEER was then measured once per day for up to two weeks. Control monolayers reached a TEER plateau 6-8 days after plating (Fig. 4i,j). However, Spef1 KD monolayers demonstrated a significant delay in the rise of TEER and eventually plateaued at a much lower resistance value [Control = 422 ± 17 Ω.cm^2^ *vs* Spef1 KD = 241 ± 24 Ω.cm^2^ TEER average values taken at 5 days of dox induction] (Fig. 4i). Additionally, Spef1 KD monolayers showed a decreased TEER measurements after three days in dox [Control = 420 ± 48 Ω.cm^2^ *vs* Spef1 KD = 163 ± 44 Ω.cm^2^] (Fig. 4j). Collectively, these results suggest that Spef1 contributes to the normal architecture of tricellular and bicellular tight junctions, and in turn, the development and maintenance of epithelial barrier function.

### Spef1 depletion leads to redistribution of non-muscle myosin-2A and increased activation

The defects in monolayer integrity and barrier function observed in Spef1 depleted cells are reminiscent of the loss-of-function phenotypes reported for p120, EpCAM, Afadin, ZO-1, Dbl3, MRCK, gp135, KIBRA and several polarity proteins (Par3/Par6/PKCz/Cdc42; Lgl/Scrib/Dlg), among others ^32–43^. One unifying feature of these phenotypes is increased actomyosin contractility. To examine the impact of Spef1 depletion on actomyosin contractility in fully differentiated Caco-2 BBE monolayers (21 DPC). We examined the localization of non-muscle myosin-2 isoforms (NM2A/C) and phosphorylated myosin regulatory light chain (p-MLC), the latter of which is an indicator of active myosin. As expected, in control monolayers, NM2C exhibited strong localization to the circumferential belt of actin that comprises the junctional complexes and punctate signal throughout the sub-apical terminal web (Fig. S2a, left). Although Spef1 depletion induced subtle changes in NM2C signal in both regions along with a marginal increase in cytoplasmic signal, the total fluorescence intensity was unchanged [Control = 223 ± 19.3 *vs* Spef1 KD = 206 ± 11.9] (Fig. S2a,d). NM2A, a non-muscle myosin-2 variant that demonstrates higher contractile activity than NM2C ^44, 45^, also localized sub-apically and in the circumferential actomyosin belt (Fig. S2b, left). However, in Spef1 KD cells, the apical levels of NM2A were increased. Quantification of total IF intensity revealed that NM2A staining was significantly higher compared to controls [Control = 345 ± 24 *vs* Spef1 KD = 846 ± 159] (Fig. S2b,e). Strikingly, Spef1 KD cells also displayed an increased p-MLC at the apical surface relative to control [Control = 192 ± 46 *vs* Spef1 KD = 380 ± 6] (Fig. S2c,f).

To follow the dynamics of NM2A accumulation, Caco-2 BBE cells were plated through cell differentiation (3 - 21 DPC), and monolayers were then immunostained for F-actin and NM2A (Fig. S3). Our findings revealed that control monolayers developed a dense array of MV structures through the course of cell polarization as demonstrated by F-actin localization (Fig. S3a, Fig. 5a, top). Conversely, Spef1 depletion attenuated MV assembly even when monolayers were fully polarized (Fig. S3a, Fig. 5a, bottom). In control conditions, NM2A was apically positioned and its distribution was gradually reduced as cells established apical-basal polarization (Fig. S3b, Fig. 5b, top). However, Spef1 loss induced a marked increase of NM2A at the junctional areas and apical surface (Fig. S3b, Fig. 5b, bottom). Quantification through the *z*-stacks from the cell-cell contacts demonstrated that NM2A significantly increased at early polarization stages and its junctional localization was sustained until cells reached the fully polarization state [Control (3 DPC) = 327 ± 116 *vs* Spef1 KD = 652 ± 69; Control (7 DPC) = 215 ± 7 *vs* Spef1 KD = 601 ± 98; Control (14 DPC) = 139 ± 38 *vs* Spef1 KD = 637 ± 97; Control (21 DPC) = 141 ± 27 *vs* Spef1 KD = 671 ± 186] (Fig. 5c-f). Together these data suggest that Spef1 disruption leads to a redistribution and increased activity of NM2A which impact the MV assembly in intestinal epithelia.

**Fig. 5:**
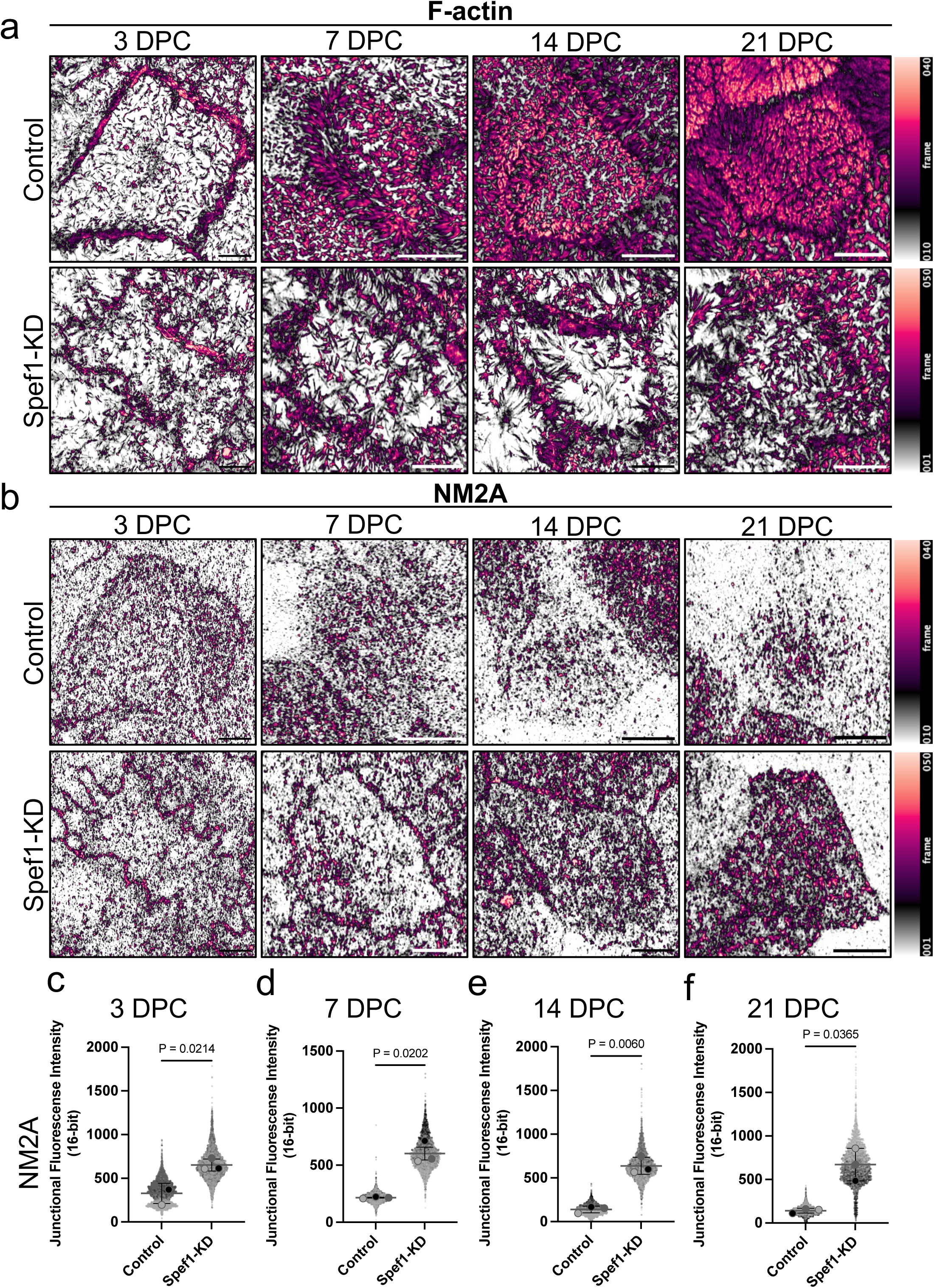
Spef1 regulates MV formation through high levels of junctional NM2A. **a,b,** Confocal projections of control and Spef1 KD cells plated at 3 - 21 DPC immunolabeled with F-actin (**a**) and NM2A (**b**) color coded. Images correspond to the dotted boxes from Fig.S3. Scale bars, 5 μm. Spef1 KD cells exhibited a delayed MV formation and increased of apical and junctional recruitment of NM2A compared with control condition. **c-f,** Superplots represent the quantification of junctional fluorescence intensity of NM2A in control and Spef1 KD monolayers through cell polarization (3 - 21 DPC). The experiments were conducted three times independently (*n* = 3). Total number of fields imaged: **c,** 3 DPC = 12; **d,** 7 DPC = 15; **e,**14 DPC = 15; **f,** 21 DPC = 18. *P* values were calculated using two-tailed *t*-test.

Our data indicate that AMIs and BACs derive from the apical membrane surface in Spef1 depleted cells. To investigate if NM2A is part of these features, Spef1 KD cells were fixed and stained for NM2A, occludin and F-actin. Confocal imaging, segmentation, and reconstruction of the resulting volumes revealed that, at early stages of Spef1 KD cell differentiation (7 DPC), NM2A and occludin were highly enriched at multicellular vertices and these signals extend down the lateral surface in space between cells at the tricellular interface (Fig. S4, top). At later stages of differentiation (14 DPC), BACs were encircled by ring-like occludin and NM2A signals remained attached to the apical surface (Fig. S4, middle). By the latest time point (21 DPC), BACs lose contact with the apical surface, but retain strong F-actin, NM2A and occludin signals (Fig. S4, bottom). Together, these data strongly suggest that Spef1 regulates the recruitment of NM2A to the apical surface and to the tricellular vertices, then impacting actomyosin contractility at these sites.

### Inhibition of non-muscle myosin-2 rescues the junctional defects caused by Spef1 depletion

To determine if AMIs induced by Spef1 depletion were associated with excess actomyosin contractility, we labeled Caco-2 BBE control and Spef1 KD cells with a membrane dye (CellBrite) and then subjected monolayers to live imaging (Fig. 6a,c). Time-lapse image volumes showed that Spef1 KD monolayers accumulated the membrane tracer at TCJs at much higher levels than control cells (Fig. 6b,d,g). Lateral views of these regions revealed that the membrane probe signal extended down into the tricellular interface reaching the basal surface (Fig. 6d, -*xz* sections). To explore whether the membrane accumulation at TCJs promoted by Spef1 depletion is due to increased actomyosin contractility, Spef1 KD cells were treated with the NM2 inhibitor, blebbistatin. Strikingly, reducing myosin contractility with blebbistatin led to a significant decrease in membrane accumulation in TCJs over time (Fig. 6e-g), suggesting that high levels of actomyosin contractility surrounding tricellular interfaces contribute to AMI formation.

**Fig. 6:**
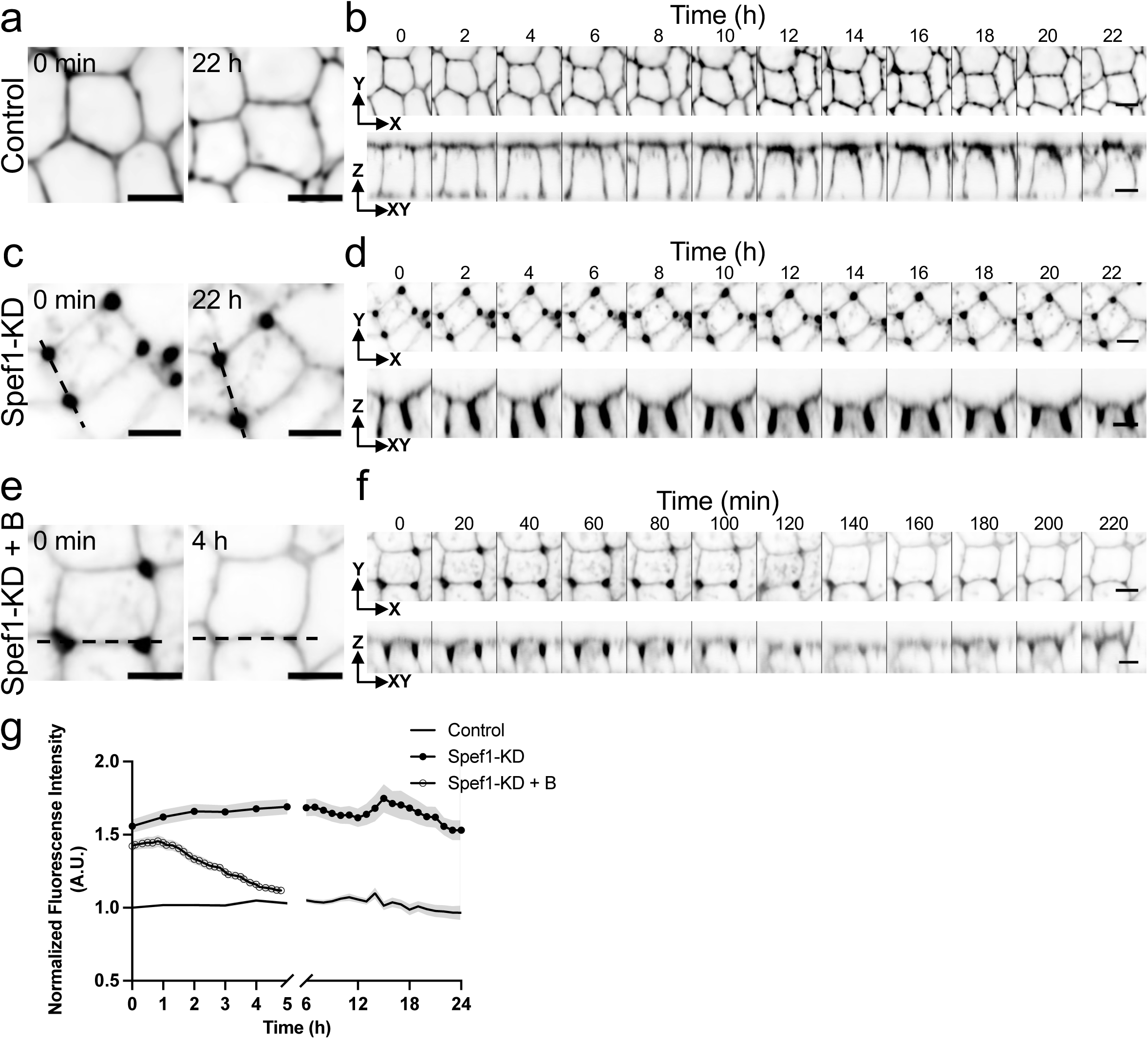
Spef1 modulates the opening of tricellular vertices through the regulation of actomyosin contractility. **a**-**d,** Confocal sections of living control (**a,b**) and Spef1 KD (**c,d**) cells incubated with the cell membrane dye, CellBrite (black). Dashed lines indicate the tricellular vertex region. Montage time-lapse imaging from control (**b**) and Spef1 KD cells (**d**) showing the top (-*xy* sections) and lateral (-*xz* sections) views indicating the tubular formation (dashed lines from **c** in Spef1-depleted cells. Images were collected every 30 min for 24 h. **e,f,** Time-lapse recording of Spef1 KD monolayers incubated with cell membrane dye (black) treated with the inhibitor of myosins, blebbistatin (B, 5 μM) for 4 h. Representative images indicate the top (-*xy* sections) and lateral (-*xz* sections) views of the small membrane invaginations of Spef1 KD cells through time. Cells were imaged every 10 min for 5 h. Scale bars, 10 μm. **g,** Quantification of the membrane tricellular accumulation through time; each data point represents the average measurements from live time-lapse images normalized regarding the control condition (control = 25; Spef1 KD = 17; Spef1 KD + Blebbistatin = 21).

## DISCUSSION

Epithelial sheet integrity is directly linked to robust adherent cell-cell contacts, which are formed and shaped using contractile forces generated by the actomyosin cytoskeleton. Previous studies established that some of these forces originate in a belt of F-actin and NM2, which encircles the cell at the interface between apical and basolateral domains ^8, 10–13, 15^. In this work, we show that in polarizing intestinal epithelial cell culture models, Spef1 loss-of-function produced striking defects in the apical domain and loss of epithelial integrity, characterized by the disruption of the TCJs and aberrant lateral accumulation of apical material. Our experiments revealed that Spef1 KD induced changes in the composition of the sub-apical compartment leading to a strong accumulation of NM2A. We propose that higher levels of contractility associated with NM2A accumulation in this region drive the aberrant junctional remodeling and tricellular contact failure observed in Spef1 KD monolayers. Below we discuss our rationale for this proposal and potential mechanisms in more detail.

Here we report that in the intestinal tract, Spef1 is highly enriched at the apical domain of villus enterocytes and differentiating cells in the crypt. Notably, this epithelial tissue does not build cilia, and thus, functions for Spef1 in this context are likely distinct from previously published roles in the axoneme ^19, 20, 25^. We found that Spef1 exhibited reduced localization at the basolateral margins of cells, and as enterocytes differentiate during migration up the crypt-villus axis, Spef1 signal shifted from basal to apical sites. These observations agree with our previous studies, which established that Spef1 localization depends on apico-basal polarization state in intestinal epithelial cells ^26^.

Using an inducible KD model, we found that Spef1 depletion led to severe defects in epithelial morphology at both cell and monolayer scales, characterized by a loss of tricellulin from tricellular contacts, and the opening of prominent gaps between cells at these sites. Tricellular contacts are sites of high tension that are essential for maintaining epithelial homeostasis ^46^. Actin crosslinker fimbrin (Taneja et al., 2025, BioRxiv unpublished data), adherens junction components (E-cad, β-catenin, vinculin, canoe/afadin), and myosin-2 motors are all enriched at tricellular vertices, where they serve to stabilize these structures and allow cells to tolerate increased levels of tension at these sites ^46–50^. Additionally, depletion of tricellular junction components such as tricellulin, lipolysis-stimulated lipoprotein receptor (LSR) or immunoglobulin-like domain-containing receptor (ILDR) 1/2 led to the formation of tricellular gaps characterized by the aberrant accumulation of ZO-1 and occludin at these sites, and compromised epithelial barrier function ^28, 29, 46, 51–53^; these phenotypes are similar to what we observed in the current study. Our data suggest that Spef1, by controlling actomyosin contractility, regulates junctional integrity and epithelia function remotely, from its apical localization in brush border MV.

How might Spef1, a resident of the microvillus, regulate NM2 contractility and in turn, junctional integrity? A microvillus is a simple structure, supported by a core of 20-30 actin filaments bundled in parallel and anchored in a dense, sub-apical filamentous network, known as the terminal web ^54, 55^. The terminal web is meshwork of intermediate filaments, actin, spectrins, and bipolar contractile filaments of NM2 which is dense enough to exclude vesicles, large organelles, and microtubules ^12, 56^.The segment of the microvillus that reaches down into the terminal web is known as the rootlet, and classic TEM images were the first to indicate that bipolar contractile filaments of NM2 make direct contact with this part of the core bundle. More recent super-resolution imaging studies of NM2C in native intestinal tissues strongly indicates that this motor localizes to the terminal web and makes connections with junctional complexes ^15^. Such physical continuity suggests that this network could propagate or generate additional tension to supplement the force produced by the junctional belt. Careful inspection of the super-resolution images in the current study revealed that Spef1 extends down along the rootlet and therefore is well-positioned to regulate NM2 binding at these sites. Loss of Spef1 induced changes in the composition of the sub-apical compartment; whereas NM2C was normally enriched in this region, Spef1 KD led to a strong accumulation of NM2A throughout the terminal web. Because NM2C and NM2A are low and high contractility myosins ^57^, respectively, these results suggested that the tension-generating potential of Spef1 KD epithelial monolayers was abnormally elevated. We propose that higher levels of contractility associated with NM2A accumulation in this region drive the aberrant junctional remodeling and tricellular contact failure observed in Spef1 KD monolayers. The integration of junctional and terminal web structures also offers a mechanism whereby perturbation of a microvillar component, in this case Spef1, might impact apical contractility and in turn, junctional integrity.

Spef1 shares homology with several cytoskeletal proteins that bind actin and microtubules including ACF7, α-parvin, nesprin, β-spectrin, plectin, MICAL, smoothelin, utrophin, EHBP1, α-actinin, filamin, dystrophin, dystonin, and EB1/3 ^18, 26^. Recent structural studies on the axoneme elucidated the residues of Spef1 responsible for binding to α− and β-tubulin (Legal et al., 2025, BioRxiv unpublished data), although the residues responsible for actin binding remain unresolved. Given the highly specific localization of Spef1 to actin bundle-supported MV reported here and the lack of cilia on the apical surface of native enterocytes, a role for axoneme/microtubule-binding in the KD phenotypes uncovered here remains unlikely. The current study reveals that Spef1 controls intestinal epithelia architecture and function by the regulating tension across the apical surface. Future studies will be required to explore the role of Spef1 in other tissues that express high levels of this factor, including lung, bronchus, nasopharynx, testis, fallopian tube, among others ^58^.

## ACKNOWLEDGMENTS

The authors would like to thank all members of the M.J.T. laboratory for their constructive feedback. We acknowledge the Loyola Core Facility Cytometry. This work was supported by the NIH grants DK125546 and DK111949 (M.J.T); DK097043 (G.H) and Veterans Affairs Merit Review Award, I01BX002687 (G.H.).

## AUTHOR CONTRIBUTIONS

Conceptualization, R.T., G.H. and M.J.T.; Methodology, R.T., A.M. and M.J.T.; Formal Analysis, R.T. and M.J.T.; Investigation, R.T. and M.J.T.; Writing, R.T. and M.J.T.; Visualization, R.T. and M.J.T.; Supervision, R.T. and M.J.T.; Funding Acquisition, M.J.T.; All authors contributed to revising the manuscript.

## METHODS

### Mouse tissue preparation

Animal studies were conducted according to National Institutes of Health and American Veterinary Medical Association guidelines by a protocol approved by the Institutional Animal Care and Use Committee, Vanderbilt University. Segments of small intestine were removed and processed following reported protocols ^15, 59^.

### Cell culture

Caco-2 BBE, SKCO-15, and Ls174T-W4 (W4) cells, were cultured in DMEM with high glucose and 2mM L-glutamine supplemented with 10% tetracycline-free fetal bovine serum (FBS) and maintained at 37°C and 5% CO_2_. For W4 cells, blasticidin (10 μg/ml), G418 (1 mg/ml), and phleomycin (20 μg/ml) were added to the media. This cell line was generous gift from Dr. Hans Clevers (Utrecht University, Netherlands).

### Lentivirus Production

HEK293FT cells (3 x 10^6^) plated on P-10 cm culture dishes were transfected with a mix containing: 1 ml of Opti-MEM (GIBCO #331985-070); 4 μg of PAX_2_ (viral packing plasmid); 2 μg pMD2.g (envelope viral plasmid); 4 μg of Spef1 KD (pInducer20-TRE-EGFP:miR30-hSpef1[shRNA#8], neomycin (G418) resistant backbone, Adgene) plasmid and 22.5 ml of 1 mg/ml PEI. The next day, media was changed, and viral particles were collected 48 - 72 h after transfection.

### Generation of Caco-2 BBE and SKCO-15 inducible system

Caco-2 BBE and SKCO-15 (3-5 x 10^5^) plated in 6-well plates were transduced with a mix of 0.5 - 1 ml of Lenti-Spef1-KD; 3.5 ml of culture media; 6-8 μg/ml Polybrene. After 48 h infection, cells were split and placed under 100 μg/ml G418 antibiotic selection for 2 weeks. Inducible Spef1 knock down was activated with 1 μg/ml doxycycline and cells were sorted for EGFP.

### Fluorescence activated cell sorting (FACS)

Caco-2 BBE and SKCO-15 Spef1 KD cells (5 - 10 x 10^6^) were pelleted and resuspended in high glucose DMEM with 20% FBS, 1X Penicillin/Streptomycin (Sigma Aldrich #P4333) and 50 μg/ml Gentamicin using 5 ml Polystyrene tubes. Cells were sorted by Loyola Core Facility Cytometry, FACSAriaII (P22300052) Cytometer, FACSDiva Version 6.1.3. EGFP positive cells were placed into a 24-well plate containing DMEM with 10% FBS and 100 μg/ml G418.

### Transfections

Transient transfections were performed using Lipofectamine 2000 (Invitrogen, #11668019,) according to the manufacturer instructions. After 16 h transfection, W4 cells were plated on coverslips and incubated in the presence of 1 μg/ml of doxycycline for 24 - 48 h to induce brush border assembly, after which cells were processed for IF.

### Tissue and cell immunolabeling

Paraffin embedded sections were deparaffinized in Histo-Clear II (National Diagnostics) two times 3 min each. Tissues were rehydrated in ethanol (100%, 100%, 95%, 90%, 70%, 50%) 5 min each, washed in PBS. Slides were incubated in antigen retrieval buffer (10 mM Tris, 0.5 mM Ethylene glycol-bis(β-aminoethyl)N,N,N′,N′-tetraacetic Acid, pH 9.0) for 1 h. Samples were rinsed three times with PBS and blocked in 10% goat serum for 1 h at RT. Primary antibody was incubated overnight at 4°C in a humidity chamber. Samples were washed with PBS and secondary antibody was incubated for 1 h at RT. Slides were rinsed three times in PBS and dehydrated with ascending ethanol series (50%, 70%, 90%,95%,100%) 5 min each. Frozen sections were thawed and rinsed with PBS to remove the OCT. Samples were permeabilized with 0.2% Triton X-100 for 10 min at RT, washed with PBS and blocked in 10% serum albumin (BSA, Research Products International) for 2 h at 37°C. Cells plated on acid-washed glass coverslips (12 mm diameter, #1.5) were fixed and permeabilized following different protocols: a) 4% PFA for 20 min at RT, washed three times with PBS and permeabilized with 0.1% Triton X-100 in PBS for 10 min at RT; b) Incubation with either cold methanol or cold ethanol at -20°C for 10 min, then cells were hydrated in PBS three times at RT. Cells were blocked with 5% BSA in PBS for 1 h at RT. Primary antibodies were diluted in 5% BSA and incubated overnight at 4°C in a humidified chamber. Cells and frozen sections were rinsed with PBS and incubated with secondary antibody for 1 and 2 h at RT, respectively. Secondary antibodies were rinsed and samples were mounted using ProLong^TM^ Gold Antifade Mountant reagent (P36930, Invitrogen).

### Antibodies

The following antibodies were used for tissue, cell staining and western blotting: anti-villin (1:50, mouse, Santa Cruz Biotechnology #SC-66022); anti-NM2A (1:100, rabbit, Biolegends #909801); anti-MLC (1:100, mouse, Cell signaling #3675); anti-NM2C (1:100, rabbit, Proteintech #20716-1-AP); anti-Spef1 (1:100 rabbit, Santa Cruz Biotechnology #SC-85485); anti-ZO-1 (1:100, rabbit, Invitrogen #61-7300); anti-occludin (1:100, mouse, Invitrogen #OC-3F10); anti-Tricellulin (1:100, rabbit, Invitrogen #488400); anti-EGFP (1:1000, chicken, Aves Labs Inc #GFP-1020); anti-GAPHD (1:1000, mouse, Santa Cruz Biotechnology #sc-32233). Secondary antibodies: Alexa Fluor 488, Alexa Fluor 568, Alexa Fluor 647, and Alexa Fluor 405-phalloidin (1:200) (Invitrogen).

### Immunoblotting

Cells were washed with ice-cold PBS, scraped and incubated on ice in lysis buffer (CellLytic^TM^ M, Sigma #C2978) containing protease inhibitors (Complete, Roche #05892791001). Lysates were boiled for 10 min in 4X Laemmli Sample buffer containing 2-mercaptoethanol (Sigma #M3148) and separated by SDS-PAGE (NuPAGE^TM^ Bis-Tris Gel NP0323BOX, Invitrogen). Gels were transferred to PVDF membranes, using a Trans-Blot Turbo Transfer System (BIO-RAD). Membranes were blocked with 5% BSA and incubated with primary antibodies overnight at 4°C, followed by three washes with PBST (1X PBS and 0.1% Tween-20). Secondary antibodies (1:5000, IRdye 800 donkey anti-rabbit, LI-COR #926-32213; IRdye 800 donkey anti-mouse, LI-COR #926–32212) were incubated for 1 h at RT in the dark. Proteins were detected by chemiluminescence using the Odyssey scanner (LI-COR). Densitometry of immunoblot data were performed using ImageJ software.

### TEER

Caco-2 BBE and SKCO-15 (3 x 10^5^) cells were seeded on the apical side of the Transwell (Costar #3470). To induce the Spef1 KD, some Transwells received 1 μg/ml of dox at day 0 (D0) DPC (TJ assembly) or at D9 DPC (TJ disassembly). TEER measurements were taken every day for two weeks using a voltohmmeter device (EVOM3, World precision Instruments). TEER values were calculated by subtracting the blank sample from each sample and multiplied for the filter area (0.33 cm^2^), values are reported as ohms.cm^2^.

### Microscope imaging

Nikon A1plus inverted microscope equipped with 405 nm, 488 nm, 568 nm and 647 nm lasers was used to perform confocal microscopy. Paraffin mouse sections were visualized using a Apo LWD 25x 1.10w DIC N2, 1.1 numerical aperture lens. Cells were imaged with Plan Apo 60x oil DIC H, 1.4 numerical aperture lens and Plan Apo TIRF 100x oil DIC H N2, 1.45 numerical aperture lens. High resolution volumes were collected using an AX/AXR confocal microscope with NSPARC, equipped with the Plan Apo λD 100x oil OFN25 DIC N2, 1.45 numerical aperture lens. All samples were imaged at 1024 x 1024 resolution, z-stacks were taken from the bottom to top with a step size of 0.1 μm. Live imaging was performed using a Spinning Disk Confocal microscope, with a 647 nm excitation laser using a Nikon Plan Apo λS 40xC SIL,1.25 numerical aperture lens. Caco-2 BBE cells (1.5 x 10^6^) were plated on 35 mm plasma cleaned glass-bottom dishes (Cellvis #D35-20-1.5-N). After 3 days growing in the presence or absence of doxycycline, cells were rinsed with PBS and incubated with CellBriteSteady650 membrane dye (Biotium #30108) in DMEM for 30 min at 37°C. Cells were imaged for 24 h at 30 min intervals. For experiments with blebbistatin, cells were imaged 10 min (T0) prior to blebbistatin addition at a final concentration of 5 μM. For control cells, DMSO was added to the media. Volumes were collected every 10 min for 5 h.

### IF intensity quantification

The signal intensities along bicellular junctions or along the MV of Caco-2 BBE and W4 cells, were measured in Fiji ImageJ and Nikon elements software, by using the measure tool that calculate the mean intensity along a line drawn along the junction. Each value was normalized by dividing each measured mean to the average non-junctional specific signal intensity.

### Segmentation analysis

Segmentation analysis involving thresholding and binarization tools were used to obtain quantitative information of the shape of the intestinal epithelia. Segmentation was applied to 3D rendered images along the *z*-axis, in some cases (apical surface) image filtering was used to reduce noise artifacts. Two independent binary classifiers, such as intensity and selection of separated objects were applied to these 3D-images. This feature allowed us to detect individual objects with different elongation, volume and sphericity, such as those that extend from the apical surface (AMIs) as well as structures at the bottom of the monolayers (BACs). Most of these structures were classified as a group within a range between 1 - 10 μm elongation for AMIs and 0.1–1 value for sphericity for BACs. Areas of low confidence were not included in the dataset in this analysis. Nikon Elements software was used to perform this analysis.

### Statistical Analysis

All data are presented as mean ± standard deviation (SD). T-test and ANOVA analysis were used to compare means. The statistical significance differences correspond to the P values ≤ 0.05. We used GraphPad Prism software to generate all the plots and statistical analysis.

