## Supplemental Figures for "Spef1 is a microvillar component that limits apical actomyosin contractility and preserves intestinal barrier function"

### Fig. S1

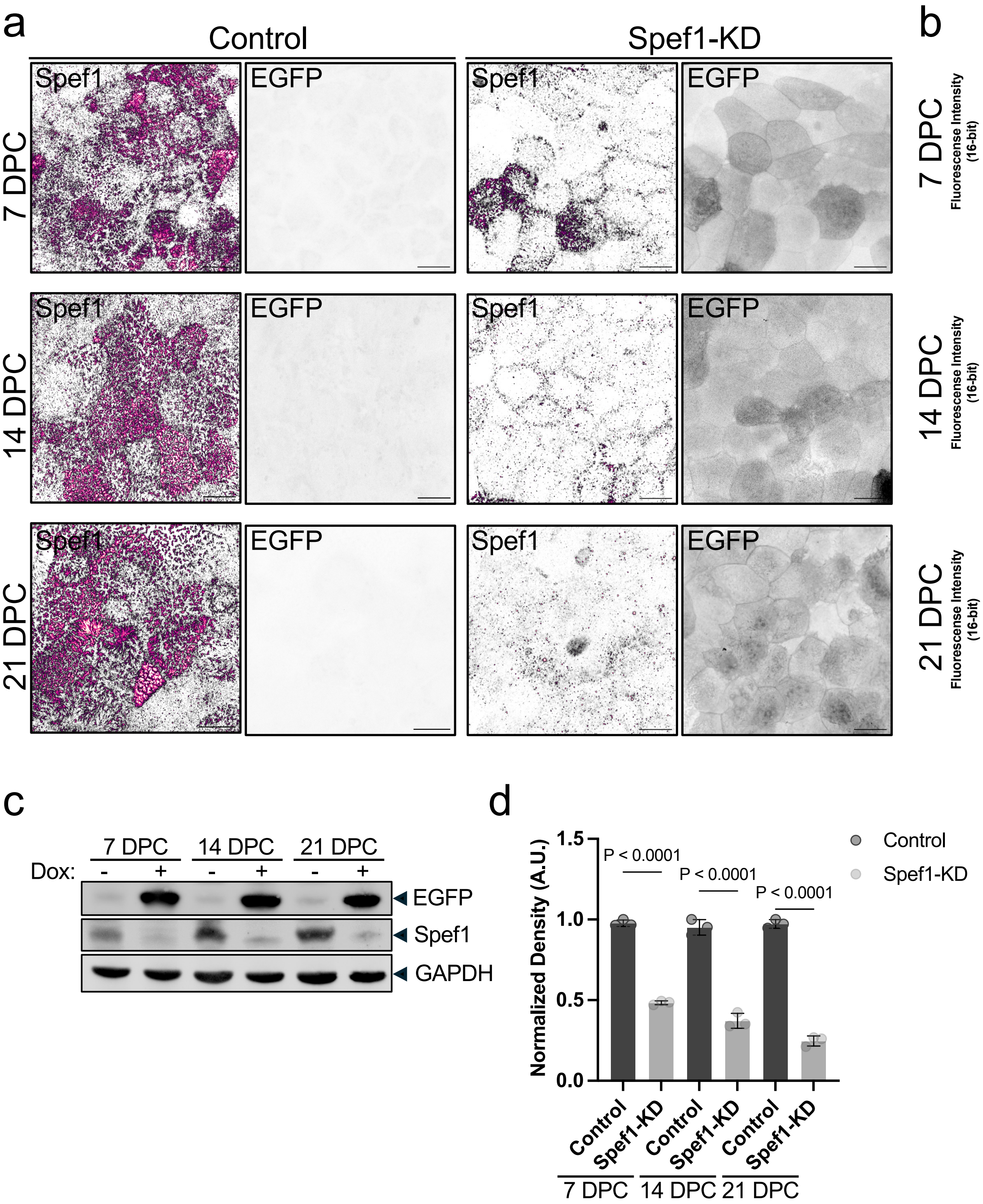
a
Control
Spef1-KD
Spef1
EGFP
7 DPC
14 DPC
21 DPC
b
P = 0.0141
P = 0.0029
P = 0.0056
Fluorescense Intensity (16-bit)
Control
Spef1-KD
c
7 DPC
14 DPC
21 DPC
Dox:
-
+
EGFP
Spef1
GAPDH
d
Normalized Density (A.U.)
P < 0.0001
Control
Spef1-KD

**Supplemental Fig.1: Generation of Spef1-KD monolayers. a,** Projected confocal volumes of Tet-On Caco-2 BBE cells in the absence (Control) or presence (Spef1-KD) of dox for 7, 14 and 21 DPC (top to bottom). Image volumes were depth-coded (grey = basal side; plum = apical side); soluble EGFP signal shown to the right indicates induction of the Tet-On system. Scale bars, 10 μm. **b,** Quantification of fluorescence intensity of endogenous Spef1 from control and Spef1 KD monolayers imaged at 7, 14 and 21 DPC (top to bottom); from three independent experiments ($n$ = 3), number of fields quantified at 7 DPC = 14; 14 DPC = 9; 21 DPC = 15. **c,** Western blotting for endogenous Spef1 in Tet-On Caco-2 BBE monolayers with or without dox grown to 7, 14, or 21 DPC. **d,** Densitometric analysis of the three independent experiments ($n$ = 3) from WB shown in **c**; EGFP and GAPDH detection levels were used as internal control of the Tet-On system and loading control, respectively. $P$ values were calculated using two-tailed $t$-test analysis for **b,** and One way ANOVA Tukeys's multiple comparisons test for **d**.

#### Fig. S2

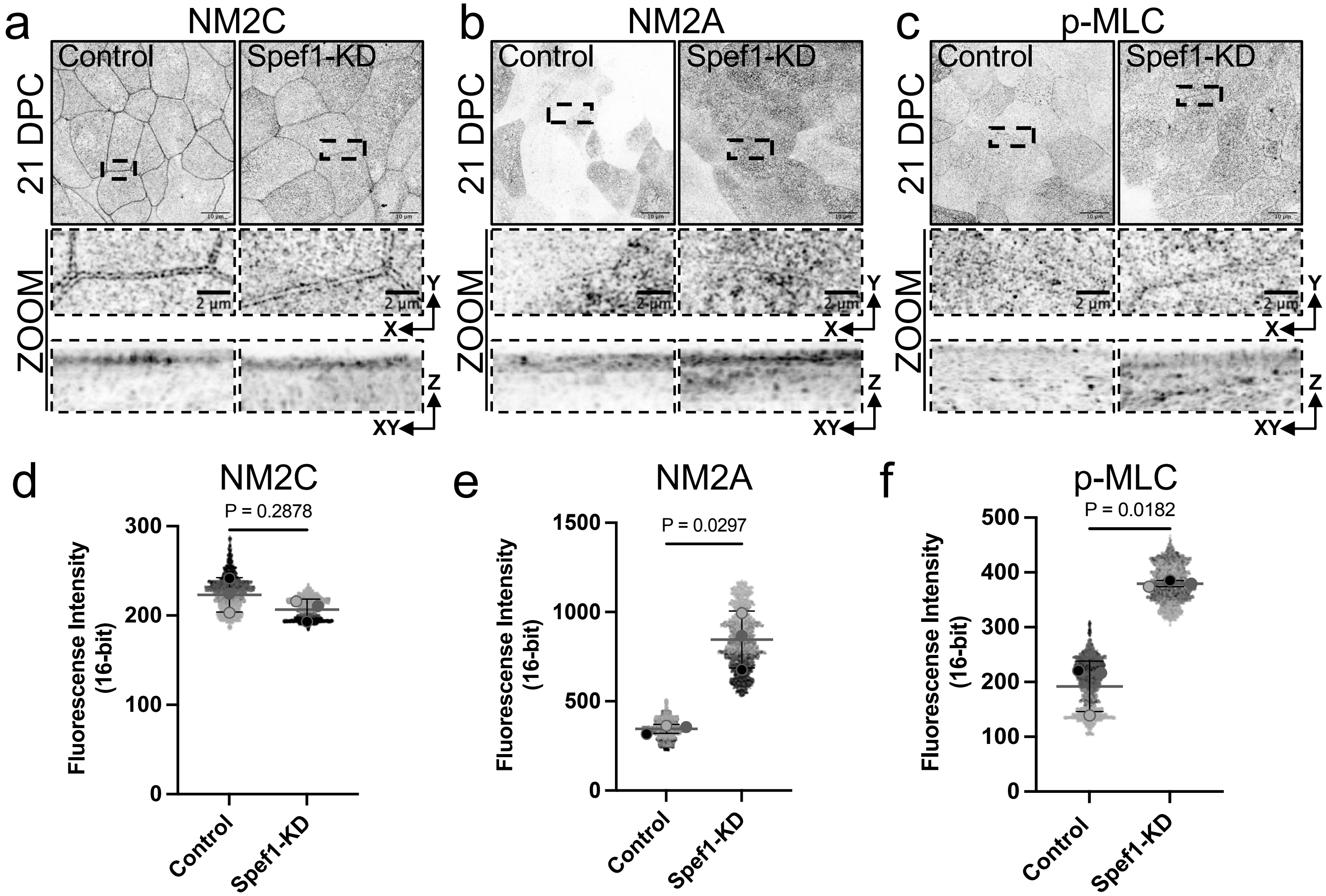
a
NM2C
b
NM2A
c
p-MLC
Control
Spef1-KD
21 DPC
ZOOM
2 µm
Y
X
Z
XY
d
NM2C
P = 0.2878
e
NM2A
P = 0.0297
f
p-MLC
P = 0.0182
Fluorescense Intensity (16-bit)
0
100
200
300
500
1000
1500
400
Control
Spef1-KD

**Supplemental Fig.2: Spef1 depletion induced high levels of NM2A and p-MLC. a-c,** Confocal images from polarized control and Spef1 KD cells (21 DPC) immunostained for NM2C (**a**); NM2A (**b**) and p-MLC (**c**). High magnification images show the localization of NM2C at the actomyosin cortex (**a,** zoom). Increased junctional recruitment and cytoplasmic accumulation of NM2A (**b,** zoom) and p-MLC (**c,** zoom) is observed in the Spef1 KD condition. **d-f,** Plots represent the average of the total fluorescence intensity of NM2C (**d**), NM2A (**e**) and p-MLC (**f**) of control and Spef1 KD monolayers. Three independent experiments ($n$ = 3) consisting of total number of fields quantified for **d:** control = 17, Spef1 KD = 19; for **e:** control = 15, Spef1 KD = 18; for **f:** control = 9, Spef1 KD = 9. $P$ values were calculated using two-tailed $t$-test.

#### Fig. S3

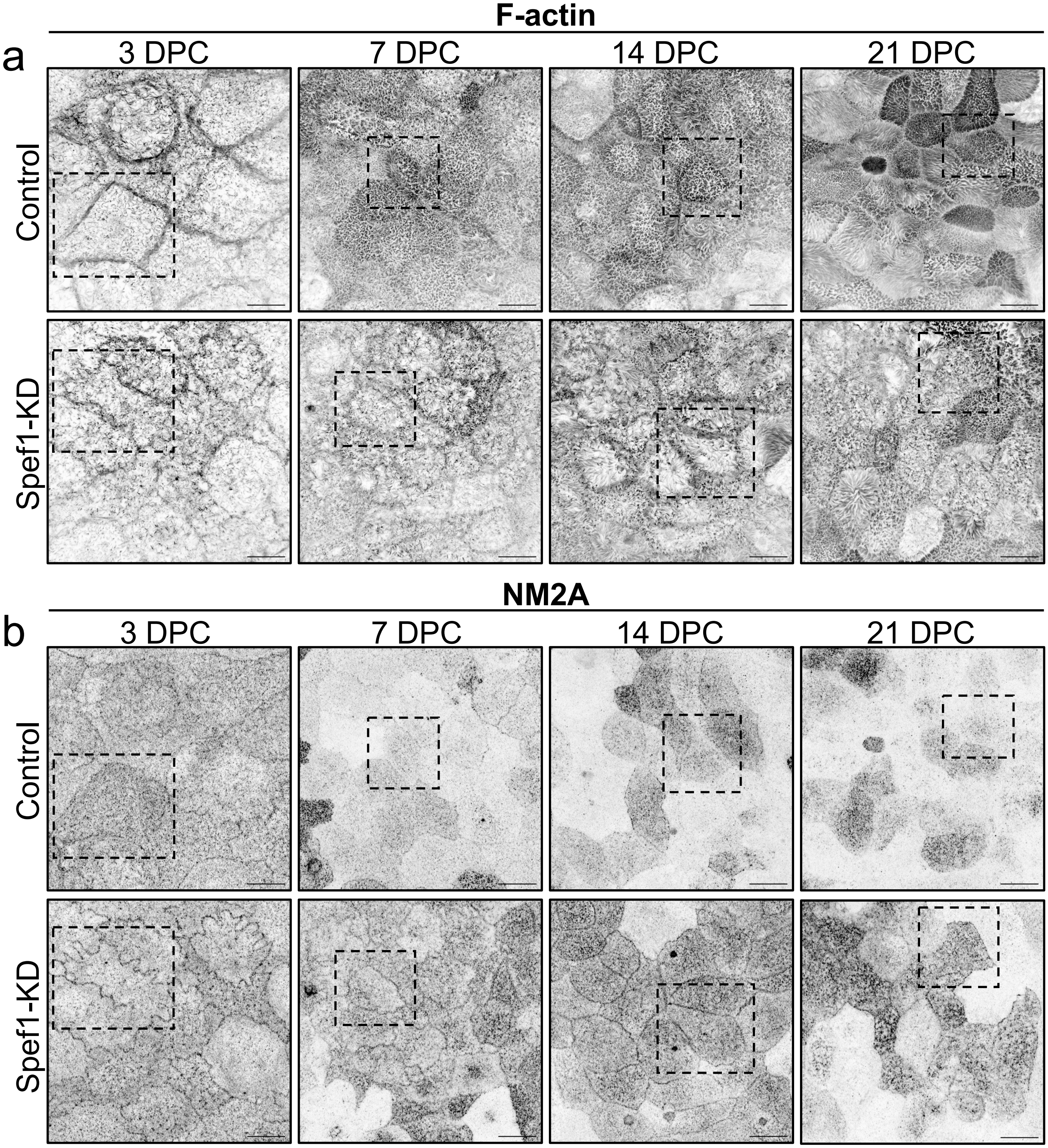
F-actin
a
3 DPC
7 DPC
14 DPC
21 DPC
Control
Spef1-KD
NM2A
b
3 DPC
7 DPC
14 DPC
21 DPC
Control
Spef1-KD

**Supplemental Fig.3: Spef1 depletion upregulates the levels of NM2A through cell polarization. a,b,** Representative confocal images of control and Spef1 KD cells plated at 3 - 21 DPC immunolabeled with F-actin (**a,** inverted channel) and NM2A (**b,** inverted channel). Scale bars, 10 μm. Spef1 KD cells exhibited a decreased MV formation and increased apical levels of NM2A compared with control condition. Total number of fields imaged per condition: 3 DPC = 12; 7 DPC = 15; 14 DPC = 15; 21 DPC = 18.

### Fig. S4

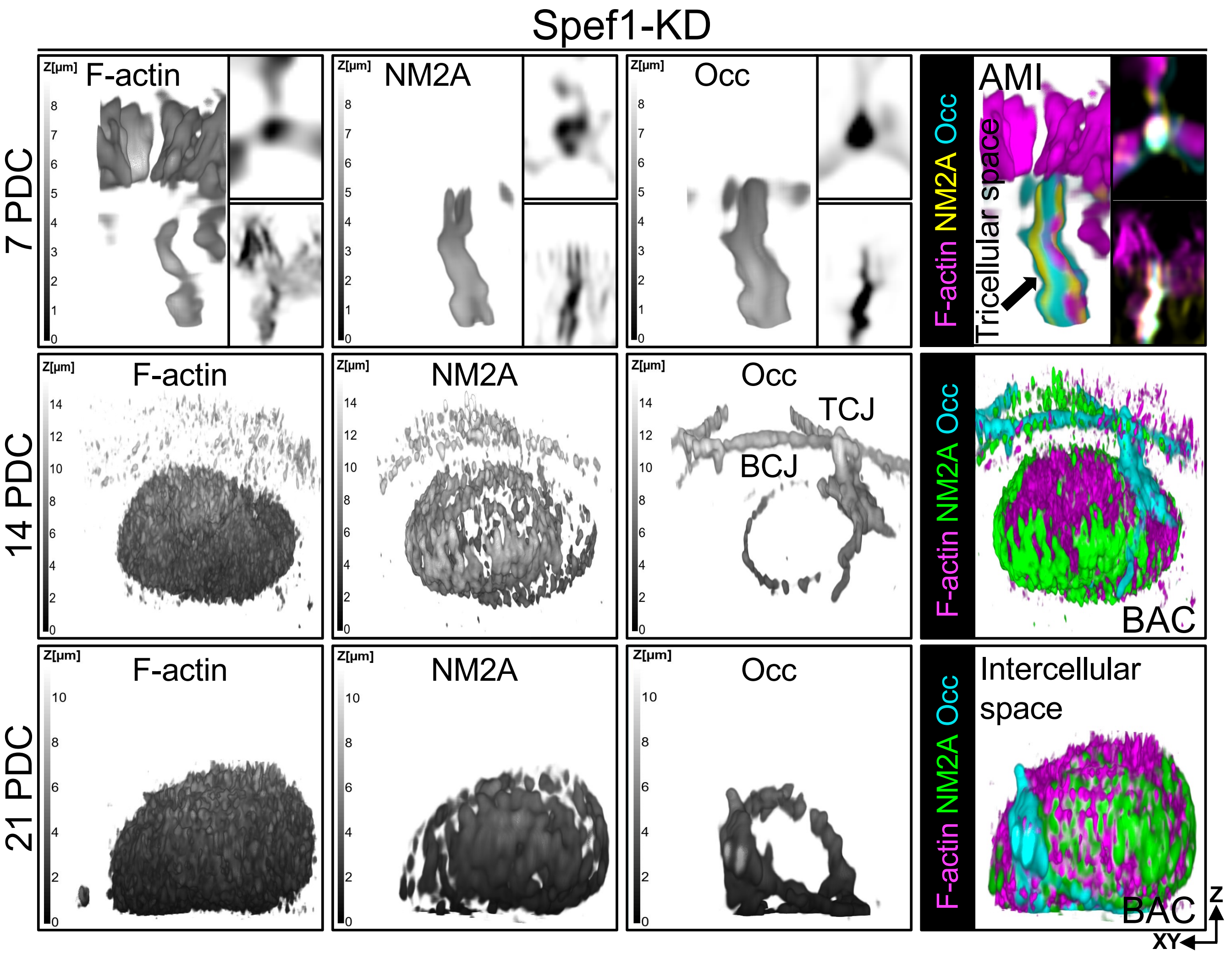

Spef1-KD
7 PDC
14 PDC
21 PDC
Z[µm]
F-actin
NM2A
Occ
AMI
Tricellular space
F-actin NM2A Occ
TCJ
BCJ
BAC
Intercellular space
Z
XY

**Supplemental Fig.4: AMIs and BACs formed during Spef1 depletion consist of adhesion and cytoskeletal molecules.** Representative 3D-rendered images from Spef1 KD cells showing AMIs and BACs in closed association with occludin and NM2A through stages of cell differentiation (7 - 21 DPC). From top to bottom, Spef1-depleted cells (7 DPC) showing F-actin (magenta), occludin (cyan) and NM2A (yellow) along (left, 3D) and enriched at the tricellular vertices (right, *-xy* and *-xz* sections). Middle, 3D-rendered images from Spef1 KD cells plated at 14 DPC showing BACs (magenta) surrounded by NM2A (green), in closed association to occludin (cyan). Bottom, 3D-rendered images from Spef1 KD cells (21 DPC) showing BACs containing occludin and NM2A proteins accumulated at the intercellular space.
